# Genome variation of Sporothrix schenckii and Sporothrix brasiliensis

**DOI:** 10.64898/2026.03.30.715206

**Authors:** Ujwal R. Bagal, Amanda R. Santos, Rodrigo Almeida Paes, Lucas Gomes de Brito Alves, Luz Rocio Chamorro, Lindsay A. Parnell, Jose Pereira Brunelli, Nancy A. Chow, Jan Pohl, Vanessa Rabello Brito, Bram Spruijtenburg, Larissa Fernandes, Bridget M. Barker, Jason E. Stajich, Maria Sueli Felipe, Johanna Rhodes, Melissa Orzechowski Xavier, Daniel R. Matute, Rosely Zancope, Marcus de Melo Teixeira

## Abstract

Sporotrichosis, a subcutaneous mycosis caused by dimorphic fungi of the *Sporothrix* genus, has become a major zoonotic epidemic in South America, primarily driven by *Sporothrix brasiliensis*. To elucidate the genomic basis of its emergence and antifungal adaptation, we analyzed whole-genome sequences from 94 *Sporothrix* isolates, integrating single-nucleotide polymorphism (SNP), copy number variation (CNV), and genome-wide association (GWAS) analyses. Comparative genomics revealed 610,242 SNPs within *S. brasiliensis* and 1,474,627 within *S. schenckii*, confirming a marked disparity in intraspecific diversity. Phylogenomic tree inference resolved six well-supported *S. brasiliensis* clades with limited internal divergence, reflecting recent population expansion, while *S. schenckii* displayed deep phylogeographic structure separating North and South American lineages. CNV profiling identified 158 affected genes in *S. brasiliensis* (60 gains, 98 losses) and 88 in *S. schenckii* (54 gains, 34 losses), concentrated near sub-telomeric regions. In *S. brasiliensis*, gains were enriched for kinases and intracellular trafficking functions, whereas losses involved genes related to translation and primary metabolism, suggesting regulatory reinforcement coupled with metabolic streamlining. A GWAS of itraconazole resistance identified 81 SNPs distributed across multiple scaffolds, with many located within genes related with transport, signaling, and redox balance, supporting a polygenic basis for azole response. Together, these results highlight distinct evolutionary strategies of closely related *Sporothrix* species and delineate the genomic changes associated with the emergence and drug tolerance of *S. brasiliensis*.

## Introduction

Fungal diseases are increasingly recognized as a major threat to human and animal health, with an estimated one billion individuals affected worldwide each year [1]. Among these, sporotrichosis has emerged as a critical and expanding public health concern, particularly in South America, where zoonotic transmission from domestic animals, especially cats, has fueled sustained outbreaks of unprecedented scale [2]. Sporotrichosis is a subcutaneous mycosis caused by thermally dimorphic fungi of the genus *Sporothrix*, with most human infections attributed to the *S. schenckii* species complex, which includes *Sporothrix schenckii* sensu stricto, *Sporothrix brasiliensis*, *Sporothrix globosa*, and *Sporothrix luriei* [3]. These fungi inhabit soil, decaying plant material, and organic debris, with transmission primarily occurring through traumatic implantation into the skin [4]. Historically, sporotrichosis was mainly a sapronotic, occupation-associated infection acquired through traumatic inoculation of *Sporothrix* spp. from soil and plant material. However, *Sporothrix brasiliensis* has emerged as a highly virulent zoonotic pathogen, driving large-scale epidemics of cat-associated sporotrichosis across Brazil, Argentina, Paraguay, Chile, and Uruguay, and has become the dominant etiological agent in South America [5]. In parallel, zoonotic transmission by *Sporothrix schenckii* has emerged in Southeast Asia and the United States, reflecting a convergent shift toward feline-associated transmission [6, 7].

The clinical manifestations of sporotrichosis vary depending on host susceptibility and fungal species. In humans, the disease typically presents as a localized cutaneous or lymphocutaneous infection, but in immunocompromised individuals, it can progress to disseminated or extracutaneous forms, with pulmonary and osteoarticular involvement [8]. In contrast, feline infections caused by *S. brasiliensis* are often severe, with high fungal burdens, extensive ulcerative lesions, and frequent systemic dissemination. Zoonotic transmission from infected cats to humans has been a major driver of the sporotrichosis epidemic in Brazil, where *S. brasiliensis* has been implicated in nearly all cases [4, 9, 10]. Unlike other *Sporothrix* species, *S. brasiliensis* isolates show enhanced virulence, higher transmissibility, and increased antifungal resistance [11, 12].

Recent genomic studies have provided insights into the evolutionary relationships among *Sporothrix* species, demonstrating that *S. brasiliensis* and *S. schenckii* are sister species, and diverged between 3.8 and 4.9 million years ago [11]. The two species differ by approximately 442,000 single-nucleotide polymorphisms (SNPs), and the dyad’s most closely related species is *S. globosa* [11, 13]. Within *S. brasiliensis*, population genomic analyses have identified several geographically restricted lineages in Brazil, suggesting diverse population structure [4, 10, 14–16]. Despite these findings, the underlying factors driving the epidemic spread of *S. brasiliensis* remain poorly understood. Specifically, little is known about how genetic variation is partitioned across the geographic range of *S. brasiliensis*. Additionally, although *S. brasiliensis* exhibits phenotypic variation such as pigmentation and growth rate, the emergence of antifungal resistance represents one of the most concerning adaptive trajectories in this pathogen [2, 17–20]. Yet, the molecular mechanisms underlying this trait remain largely unexplored (but see [12, 21]).

In this study, we applied whole-genome sequencing (WGS) to investigate intraspecific genetic variation, population structure, and large-scale structural variation, including copy number variations (CNVs), in clinical isolates of *Sporothrix schenckii* and *S. brasiliensis*. By integrating antifungal susceptibility testing with comparative genomic and CNV analyses, we aimed to identify genomic features associated with antifungal resistance, pathogenicity, and host adaptation. Our structural analyses reveal species and lineage-specific CNV landscapes, marked by contrasting patterns of gene gains and losses, heterogeneous chromosomal blocks of amplification and deletion, and distinct population-level CNV signatures. These genomic architectures highlight fundamentally different modes of genome plasticity between *S. brasiliensis* and *S. schenckii*. Given the increasing burden of sporotrichosis, particularly in South America, our findings provide critical insights into the genomic mechanisms underlying the emergence and diversification of *S. brasiliensis*, with direct implications for public health surveillance, antifungal treatment strategies, and the evolutionary dynamics of pathogenicity.

## Methods

### Genome Sequencing

We sequenced 13 new *Sporothrix* isolates, ten strains of which were isolated from Brazil and three from Paraguay. All isolates were cultured in Brain Heart Infusion (BHI) broth (Becton Dickinson and Company, MD, USA) at 37°C for seven days. Following incubation, genomic DNA was extracted from the yeast-phase cells using a previously established protocol [22]. DNA integrity was assessed by electrophoresis on 1% agarose gel with a DNA mass ladder, and by spectrophotometric analysis using a NanoDrop™ 2000 (Thermo Scientific).

We used these DNA extractions to sequence the genome of each isolate. Sequencing libraries were prepared from 1 µg of high-quality genomic DNA per sample using the KAPA HyperPrep Kit (Roche, Basel, Switzerland), following the manufacturers protocol optimized for Illumina® sequencing platforms. Each library was uniquely indexed with 8 bp nucleotide barcodes, and final concentrations were quantified using the KAPA Library Quantification Kit (Kapa Biosystems) on a 7900HT real-time PCR system (Life Technologies). Libraries were then paired-end sequenced with a read length of 250 bp, using v4 chemistry on the Illumina NovaSeq 6000 system (Illumina, San Diego, CA).

### Genomic Data

We also incorporated previously published genomic data from *S. brasiliensis* and *S. schenckii* [11, 13, 23]. **Table S1** lists all the isolates used in the study, along with their geographic origin and accession numbers. In total, we included 83 previously sequenced *Sporothrix* genomes - 75 *S. brasiliensis* isolates and 8 *S. schenckii* isolates*. The S. brasiliensis* samples included both human (69%) and animal (30%) cases of sporotrichosis. All *S. schenckii* isolates were retrieved from human samples. The isolates were collected over a span of 24 years (from 1998 to 2022), although nine samples lacked specific collection date information. Of these, 73 isolates were from Brazil and their genomic sequences were retrieved from the NCBI SRA database (**Table S1**). The 8 previously sequenced *S. schenckii* isolates originated from Brazil, Colombia and the USA.

### Data Preprocessing, Read Mapping, and Variant Calling

All FASTQ files were processed using the North Arizona SNP pipeline (NASP v1.0) [24]. NASP uses trimmomatic (v0.35) [25] for raw reads trimming, Samtools (v1.2) [26] for reference genome indexing, NUCmer aligner from MUMmer (v4) [27, 28] to identify duplicate regions and BWA-MEM (v0.7.7)[29] for mapping reads to the reference genome. Sequencing reads from each species were aligned against their respective reference genomes, *S. brasiliensis* (GCF_000820605.1) and *S. schenckii* (GCA_000961545.1). The final dataset comprised a total of 95 isolates. Next, we used the resulting BAM files to call variant sites using GATK (v3.6)[30]. We first applied the realignerTargetCreator and IndelRealigner tool from GATK to identify and generate a target interval list for local realignment. The resulting realigned BAM files were then used as input for the UnifiedGenotyper, specifying haploid ploidy and an estimated heterozygosity value of 0.01. Following variant calling, we applied the GATK VariantFiltration tool using the following filters and thresholds: Quality by Depth (QD) ≥ 2.0, Fisher Strand Bias (FS) ≥ 60.0, Mapping Quality (MQ) ≥ 30.0, MQRankSum ≥ −12.5 and ReadPosRankSum filter ≥ −8. Additionally, we excluded SNPs with < 10X coverage, < 90% variant allele frequency, and those SNPs that fell within putative duplication regions as identified by NUCmer.

The mating-type idiomorphs of both *Sporothrix* genomes were predicted by mapping sequencing reads against the full-length nucleotide sequences of the MAT1-1 and MAT1-2 loci (see [10]) using the Minimap2 tool [31] and default parameters.

### Phylogenomics and Population Genetics

Maximum likelihood (ML) phylogenetic trees were constructed using IQ-TREE2 [32], to investigate the genealogical relationships among the *Sporothrix* isolates. Within IQ-TREE2, the ModelFinder [33] feature was used to determine the optimal nucleotide substitution model by employing the TEST function to systematically evaluate and select the best-fit evolutionary model based on the Bayesian Information Criterion (BIC). Phylogenies were inferred from whole-genome single nucleotide polymorphisms (SNPs) as well as from individual scaffolds derived from *S. brasiliensis* reference genomes to evaluate genealogical concordance among genome segments. Branch support was estimated using three complementary approaches: ultrafast bootstrap approximation (UFBoot) [34] and approximate likelihood-ratio tests (aLRT)[35], each performed using 1,000 replicates and the Bayesian-like transformation of aLRT (aBayes) metric to provide reliable confidence estimates for tree topology [36]. All resulting trees were visualized using FigTree v1.4.2 (http://tree.bio.ed.ac.uk/software/figtree/).

The population structure was evaluated using two complementary population genetic approaches: principal component analysis (PCA) and model-based clustering. To further assess clustering, fastStructure v1.0 [37] and ADMIXTURE v1.3.0 [38], were used to estimate the most likely number of genetic clusters represented in the isolate dataset. For these analyses, genomic data were formatted using Plink1 [39] to generate the required input files, including .bed (binary genotype data with individual sample identifiers), .bim (SNP marker positions and annotations), .fam (sample metadata), .map (genomic positions of SNPs), and .ped (pedigree and genotype data). The number of populations (K) was evaluated across a range from 1 to 12, and the ChooseK.py script from fastStructure was used to determine the most appropriate number of clusters.

Deviation from the expected 1:1 MAT1-1: MAT1-2 ratio was evaluated independently for each clade. Because several clades contained small numbers of isolates, two-sided exact binomial tests were used as the primary inference framework, with chi-square goodness-of-fit tests reported for comparison when expected counts permitted. Clades with fewer than four isolates were interpreted cautiously, and lack of statistical significance in these cases was attributed to limited power. Associations between phylogenetic distribution and epidemiological variables (source, state, and year) were assessed, in which each clade was contrasted against all other isolates. For each variable, contingency tables were constructed after excluding isolates with unknown values. To accommodate sparse tables and low expected counts, statistical significance was evaluated using permutation-based chi-square tests of independence, in which clade labels were randomly permuted to generate empirical null distributions of the chi-square statistic. All tests were two-sided, and results are reported as permutation p-values.

### Copy Number Variation (CNV) Analysis

Initially, each assembly was split into one FASTA file per contig using Python 3.11 and Biopython 1.81. Contig headers were sanitizing to filesystem-safe names, sequences wrapped at 60 bp, and an index (index_contigs.csv) was generated that included the original headers and lengths. To scan for telomeres, the last 10 kb at both ends of each contig were analyzed for the canonical fungal hexamer TTAGGG (and its complementary CCCTAA) using TIDK [40]. The total number of overlapping motif hits and the longest exact tandem run (in base pairs and 6-mer “units”) were recorded. A contig end was considered telomeric if it contained ≥5 exact units and ≥10 motif hits, and a contigs with telomeric ends on both sides were labeled as telomer-to-telomere (T2T). Centromere candidates were inferred using a regex-based search for simple tandem arrays (repeat unit k=2–10 bp, repeated ≥12 times), merging overlaps and annotating coordinates, length, repeat unit/k, distance to each contig end, and local GC% within a ±5 kb window. The top candidate per contig was flagged as the longest array located ≥20 kb from both ends. Repeats were inferred and masked using RepeatMasker and EDTA [41].

To detect CNVs associated with genomic alterations such as gains and losses, Control-FREEC, a read depth-based approach, applied to the realigned BAM files [42]. The following parameters were used: breakpoint threshold = 0.8, ploidy = 1, minimum expected GC content = 0.33, maximum expected GC content = 0.63, read count threshold = 10, telocentromeric value = 0, and a window size of 1,000 base pairs. For each sample, the mean and variance of copy numbers were computed using a 250-bp window to quantify variation in sequencing depth along assembled genomes. To quantify the extent of CNV differentiation between populations, VST (Variance Standardized by T) was calculated, ranging from 0 (no differentiation in CN allele frequencies between populations) to 1 (complete differentiation in CN allele frequencies between populations). CNV content among lineages was compared pairwise using Wilcoxon Prank-Sum and Kolmogoroc-Smirnov Test. To visualize CNV patterns, chromosome-wide CNV heatmaps were generated using the Pandas library within an IPython environment ( Python v3.9.13) [43], showing regions of copy number loss and duplication across individual samples.VST values were also plotted along genomic positions to highlight regions of significant divergence.

### Genome-wide association analysis to identify loci associated with resistance and population structure

The R package ‘treeWAS’ (version 1.1) was used to identify loci significantly associated with antifungal drug resistance and population structure/lineage. A maximum likelihood phylogeny was constructed for isolates with known itraconazole minimum inhibitory concentrations (MICs), comprising 66 isolates with 820,230 positions as previously described. Analyses were performed in R version 4.2.3 using a binary phenotype, with an MIC cutoff of 16 to define resistance and a *p-value* cutoff of 0.01 for all three association tests. Significant *p-values* were averaged across the three tests. The same approach was repeated to identify loci significantly associated with population structure, using lineage membership provided as a discrete variable for the phenotype.

## Results

### Phylogenetic assessments

A total of 610,242 single nucleotide polymorphisms (SNPs) were identified in the *S. brasiliensis* sample (85 isolates), while *S. schenckii* (10 isolates) showed greater diversity with 1,474,627 SNPs. These variant sites were used to generate a maximum likelihood (ML) tree. As expected, the tree showed clear differentiation between the two species, demonstrating strong branch support and reciprocally monophyletic lineages (**Fig 1A**). The *S. brasiliensis* tree partition revealed six highly supported clades, each containing closely related isolates. Notably, each of these clades were geographically restricted, indicating divergence was influenced by distance within Brazil. Population genetic assessments yielded similar results. In *S. brasiliensis*, PCA revealed that a substantial portion of the genetic variation was captured by the first principal component (PC1), which accounted for 42% of the total variance (**Fig 1B**). PC1 discriminates between Clades A, B, C, D and F. PC2, which explains 24% of the variance, distinguished Clades D and E from the remaining isolates. FastStructure analysis also indicated that the genetic variance in S*. brasiliensis* is best explained by six clusters (**Fig 1C, Table S2**).

**Figure 1.**
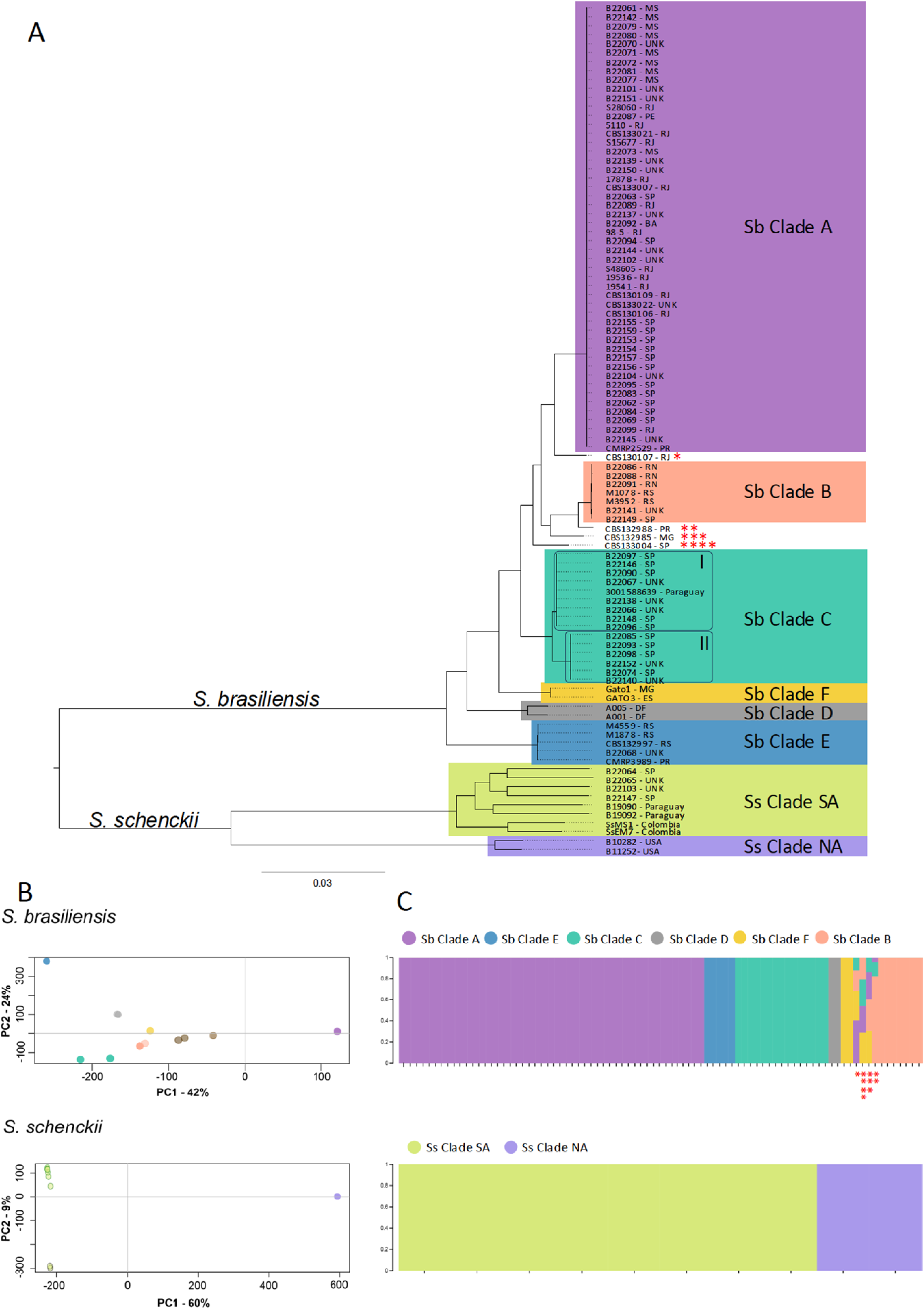
Phylogenomic structure and population stratification of *Sporothrix brasiliensis* and *S. schenckii.* (A) Maximum-likelihood (ML) phylogeny inferred from 2,274,664 genome-wide SNPs showing a strong separation between *S. brasiliensis* and *S. schenckii* isolates. Six well-supported clades were recovered within *S. brasiliensis* (A–F), whereas *S. schenckii* segregated into two major phylogeographic lineages corresponding to South America (SA) and North America (NA). Node supports are based on ultrafast bootstrap (UFBoot) and aBayes values from IQ-TREE2. (B) Principal component analysis (PCA) of 94 isolates, illustrating clustering according to species and population. *S. brasiliensis* clades cluster into discrete genetic groups, while *S. schenckii* shows clear separation between North and South American isolates. (C) Ancestry inference analysis using FastStructure and ADMIXTURE showing K = 6 populations in *S. brasiliensis* and K = 2 in *S. schenckii*. Each vertical bar represents one genome, and colors correspond to ancestral population membership. Four *S. brasiliensis* isolates (CBS130107, CBS132988, CBS132985, and CBS133004) exhibit mixed ancestry, consistent with admixture or incomplete lineage sorting

Sb Clade A, includes isolates from Rio de Janeiro, São Paulo, Pernambuco, Paraná, Bahia and Mato Grosso do Sul and is dominant. This clade was previously classified as the RJ population or H1 based on MLST data [10, 14]. It is comprised of samples from both human (n=36) and cat (n=14), with the majority collected from Brazil between 1998-2021, coinciding with the sporotrichosis epidemics. Sb Clade B (7 isolates), which includes isolates from Rio Grande do Norte, São Paulo, and Rio Grande do Sul (designated as clade RS or H3), emerged no earlier than 2012 [10, 14]. Sb Clade C is predominantly composed of human-derived isolates from São Paulo, with a single veterinary isolate (from a cat) identified in Paraguay. Two strongly supported subclades are observed within this clade. The cat was brought to Paraguay by its owners from Guarulhos, SP. The Sb Clade D consists of veterinary-derived isolates from the Federal District of Brazil, consistent with MLST (H6-7) and previous genome-typing findings [5, 10, 14]. Sb Clade E contains isolates from Rio Grande do Sul and Paraná, constituting a second cryptic *S. brasiliensis* lineage emerging in southern Brazil. Sb Clade F represents a unique genetic southeastern Brazilian lineage composed of two cat-derived isolates from Minas Gerais and Espírito Santo and previously genotyped as H2 via MLST data [14]. Isolates CBS130107 (RJ), CBS132988 (PR), CBS132985 (MG), CBS133004(SP), are positioned between strongly supported clades, a pattern seen in individual contig genealogies (**Fig S1**) which may reflect unsampled intermediate lineages, or potential cases of admixture or incomplete lineage sorting (**Fig 1A**).

In the case of *S. schenckii*, the phylogenetic tree (**Fig 1A**), PCA **(Fig 1B)** and FastStructure analyses **(Fig 1C, Table S2)** indicated the presence of two populations, one restricted to North America and the other to South America. While PC1 accounted for 60% of the genetic variation, clearly separating the South American (Ss Clade SA) and the North American (Ss Clade NA) groups, distinguishing isolates by continental origin (**Fig 1B**). PC2, which accounted for 9% of the variance, captured finer-scale genomic differentiation among South American isolates, including those from Brazil, Paraguay, and Colombia. Together, these axes reveal a strong pattern of population structure and geographic isolation within each of the two species of *Sporothrix* included in this study.

### Distribution of epidemiological variables across clades

Mating-type frequencies were evaluated within each clade against a 1:1 expectation using both exact binomial and chi-square goodness-of-fit tests. All clades were fixed for a single mating type, except Clade D (from Federal District), which was the only population containing both MAT1-1 and MAT1-2 (1:1) and therefore showed no deviation from the expected ratio (binomial p = 1.0; χ² p = 1.0). In contrast, Clade A was completely MAT1-2 (49/49) and showed the strongest distortion (binomial p = 3.55×10⁻¹⁵; χ² p = 2.56×10⁻¹²), while Clade CI and Clade B were entirely MAT1-1 (10/10 and 7/7, respectively) and Clade CII entirely MAT1-2 (6/6), consistent with significant departures from 1:1 by both frameworks (exact p ≤ 3.13×10⁻²). It’s worth nothing that Sb Clade C-I is primarily composed of MAT1-1 idiomorph, whereas Sb Clade C-II is populated by the MAT1-2 allele, suggesting two independent clonal expansions that occurred between 2017 and 2021 within São Paulo state of Brazil. Clades E and F were both fixed for MAT1-1 but did not show statistically significant deviation from the expected 1:1 mating-type due to low sample size. Collectively, these results indicate pervasive mating-type fixation across clades, with Clade D as the only exception retaining both mating types **(Table 1)**.

**Table 1.** Distribution of mating-type idiomorphs (MAT1-1 and MAT1-2) across *Sporothrix brasiliensis* clades. For each clade, the number of isolates (n) carrying MAT1-1 or MAT1-2 is shown. Deviations from the expected 1:1 mating-type ratio were tested using binomial and χ² goodness-of-fit tests.

| Clade | n | MAT1-1 | MAT1-2 | p (binomial) | $\chi^2$ | p ($\chi^2$) | min expected |
| --- | --- | --- | --- | --- | --- | --- | --- |
| Clade_A | 49 | 0 | 49 | 3.55e-15 | 49.00 | 2.56e-12 | 24.5 |
| Clade_B | 7 | 7 | 0 | 1.56e-02 | 7.00 | 8.15e-03 | 3.5 |
| Clade_CI | 10 | 10 | 0 | 1.95e-03 | 10.00 | 1.57e-03 | 5.0 |
| Clade_CII | 6 | 0 | 6 | 3.13e-02 | 6.00 | 1.43e-02 | 3.0 |
| Clade_D | 2 | 1 | 1 | 1.00 | 0.00 | 1.00 | 1.0 |
| Clade_E | 5 | 5 | 0 | 6.25e-02 | 5.00 | 2.53e-02 | 2.5 |
| Clade_F | 2 | 2 | 0 | 5.00e-01 | 2.00 | 1.57e-01 | 1.0 |

Associations between genetic clades and epidemiological variables were evaluated by testing clade-specific distributions of source, geographic origin, and sampling year against the overall dataset after exclusion of admixed isolates. To accommodate sparse data and unequal sampling, statistical significance was assessed using permutation-based chi-square tests, and inference was restricted to clades represented by at least four isolates. Under this framework, geographic state of origin showed consistent evidence of association with clade structure. In particular, Clades A, B, and E exhibited non-random state distributions relative to the global isolate set (permutation *p* = 2.0 × 10⁻⁴, 1.0 × 10⁻³, and 0.012, respectively), indicating pronounced geographic structuring. In contrast, no significant associations were detected with sampling year (all permutation *p* ≥ 0.24) or source (all permutation *p* ≥ 0.31) among clades meeting the minimum sample-size threshold. Together, these results indicate that geographic origin is the dominant epidemiological correlate of clade structure, whereas host/source and temporal variables do not show detectable clade-specific enrichment in the present dataset.

### Genome-wide CNV architecture in *S. brasiliensis* and *S. schenckii*

Structural variation within *S. brasiliensis and S. schenckii* were investigated. Telomeric ends and centromeres were identified in both genomes (GCF_000820605.1 and GCA_000961545.1). For downstream analysis of CNV, the distributions of telomeric run units and candidate centromere lengths were masked. The genome-wide CNV profiles of *S. brasiliensis* and *S. schenckii* revealed a predominant trend toward genomic losses over gains in both species, *S. schenckii* (**Fig S2**) and *S. brasiliensis* (**Fig S3**). Both species showed heterogeneous CNV landscapes that differed between them. In *S. brasiliensis*, CNVs formed well-delimited blocks alternating between amplification and loss across hundreds of kilobases (**Fig 2A**). The largest variation was observed on contig NW_024467135.1, where duplications and deletions frequently flanked sub-telomeric regions enriched in repetitive sequences. High VST (Variance based FST) values indicated regions where CNVs varied between populations. Contigs NW_024467133.1, and NW_024467134.1 displayed duplication hotspots that harbored a large portion of the detected duplications (**Fig 2A**). Overall gene losses were predominant, suggesting with large-scale deletions or pseudogenization rather than tandem expansion in *S. brasiliensis*.

**Figure 2.**
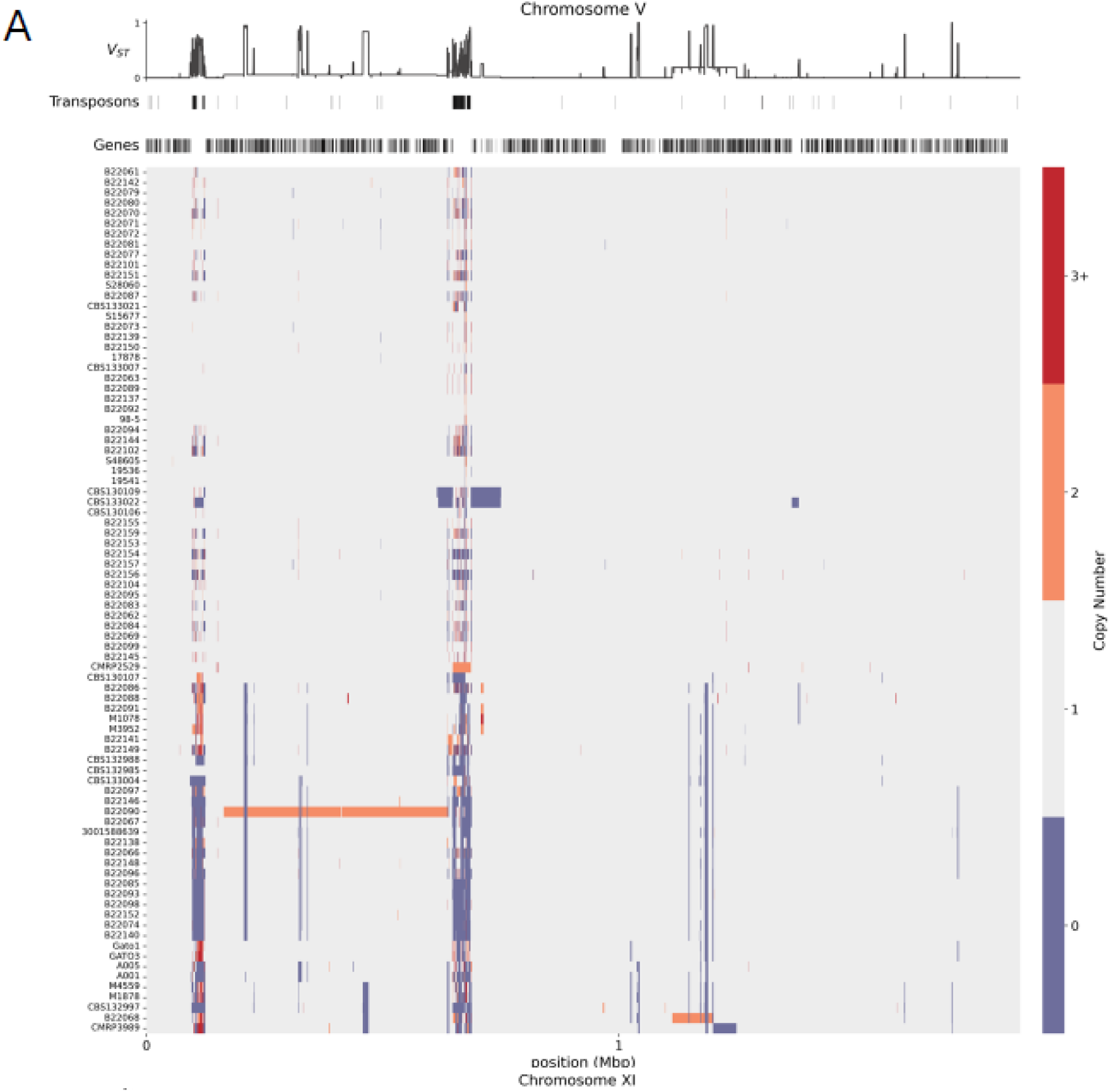
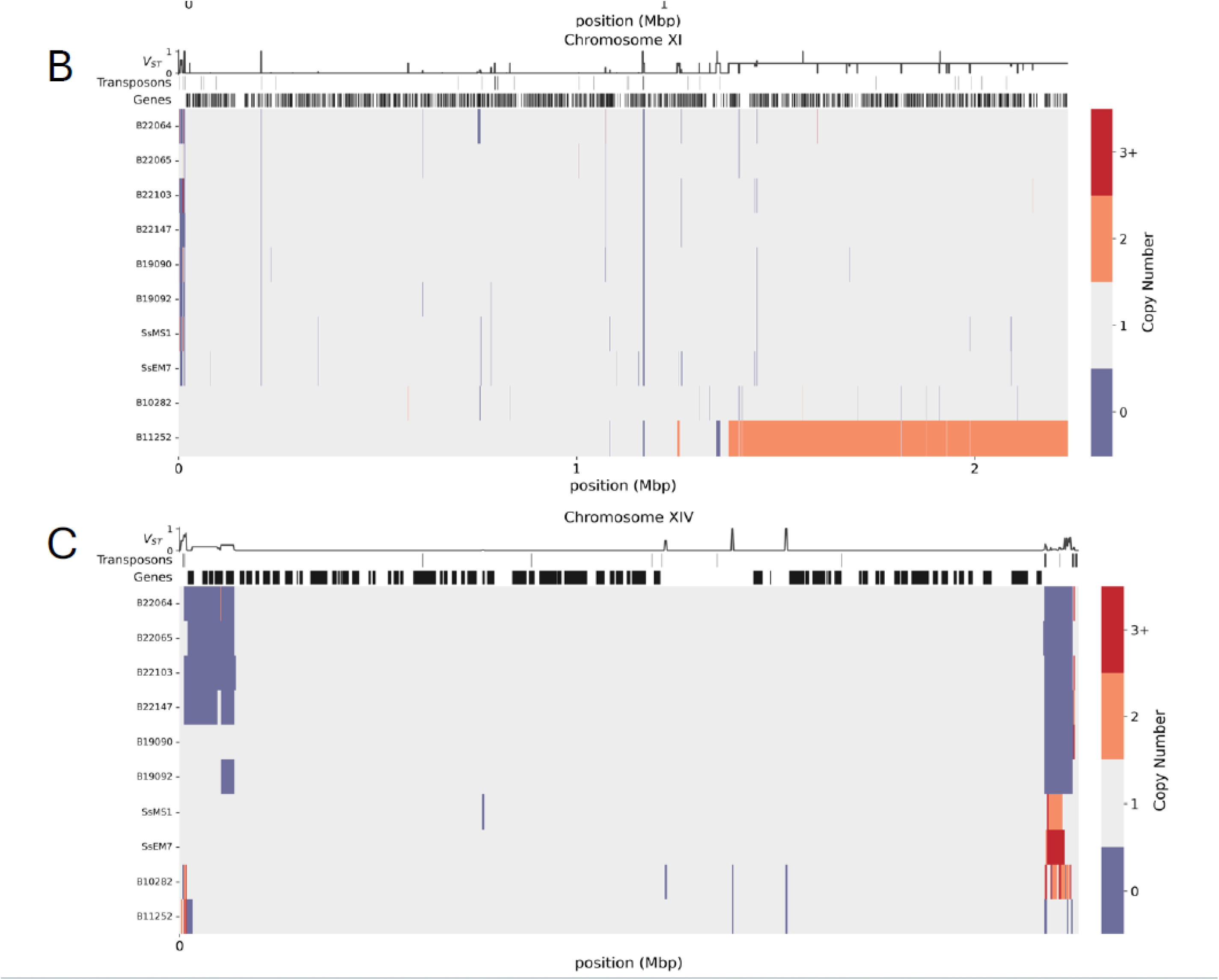
Copy number variation across representative contigs of *Sporothrix brasiliensis* and *S. schenckii.* (A) *S. brasiliensis* contig NW_024467135.1 showing large CNV blocks with alternating loss (blue) and duplication (red) events. (B) *S. schenckii* contig AXCR01000011.1, exhibiting a continuous high-amplitude duplication region exceeding 100 kb. (C) *S. schenckii* contig AXCR01000014.1, marked by irregular interspersed duplication and deletion segments generating a complex CNV landscape.

In contrast, the CNV landscape *S. schenckii* was characterized by multiple interspersed duplications and deletions (Fig. 2 B–C). Contig AXCR01000011.1 showed a large duplication block exceeding 100 kb, particularly prominent in strain B11252, and was interrupted by sharp deletion valleys resulting in a mosaic pattern of gain and loss. Contig AXCR01000014.1 displayed loss events that were polymorphic among strains (B22064, B22065, B22103, B22147, B19092 and B11252). Genome-wide, the *S. schenckii* genome had 36 CNV hotspots, predominantly concentrated on AXCR01000011.1 (16 hotspot windows) and AXCR01000006.1 (6 windows), with additional focal peaks observed on contigs AXCR01000001.1, AXCR 01000007.1, AXCR 01000010.1, AXCR 01000012.1, and AXCR 01000014.1.

Next, the gene content of the identified CNVs was characterized. A total of 158 unique CNV-affected genes were identified in *S. brasiliensis* (60 gains and 98 losses), which corresponds to approximately 1.7 % of the total predicted gene content of the reference genome (strain 5110). In *S. schenckii*, 88 genes (54 gains and 34 losses) were affected, representing about 0.5 % of the reference genome (strain 1099-18). The extent of structural variation not only differed between species but also varied within each species (**Fig 3**).

**Figure 3.**
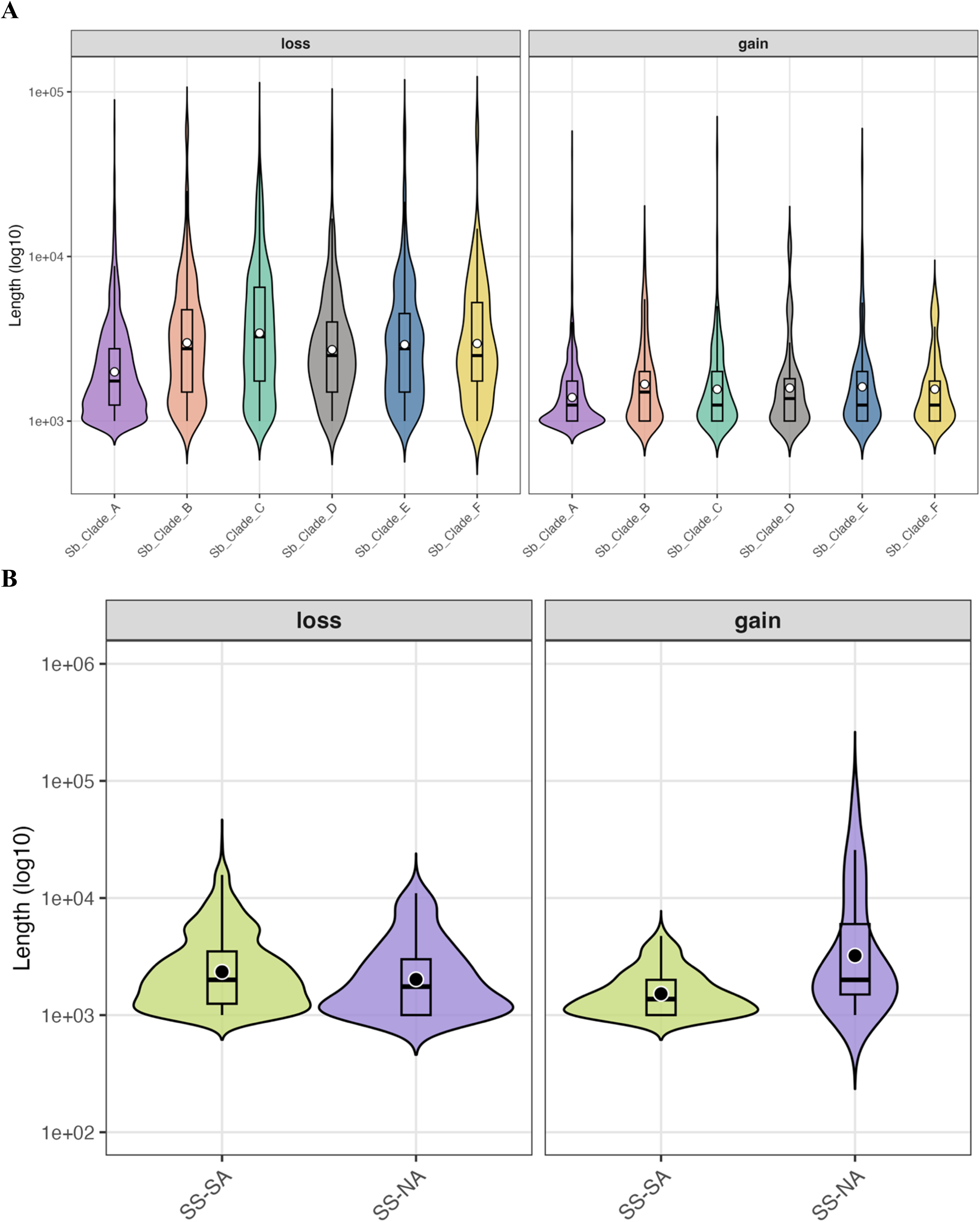
Copy Number Variation (CNV) Analysis in Sporothrix species. A Violin plot illustrating distribution of copy number variations characterized by loss and gain across various clades in A) Sporothrix brasiliensis (Sb_Clade_A, Sb_Clade_B, Sb_Clade_C, Sb_Clade_D, Sb_Clade_E, and Sb_Clade_F) and B) *S.schenckii* (South American Clade, North American Clade). The y-axis represents the length of CNVs on a logarithmic scale (log10), indicating the magnitude of the variations. The x-axis categorizes the data into different clades, allowing for a comparison of CNV patterns among them.

The CNV burden displayed considerable variability within both species, following distinct population-level patterns. In *S. brasiliensis*, most CNVs were isolate-specific, with minimal overlap between individuals. Clade Sb A carried the largest number of CNVs relative to the reference genome (22 gains, 63 losses), followed by Clade Sb C (20 gains, 41 losses) and Clade Sb E (9 gains, 34 losses). Clade F showed the lowest count, with only four loss events (**Figure 3A**). In *S. schenckii*, CNV dynamics differed between the two lineages. North American (NA) isolates showed more gains than losses relative to the reference genome (54 gains vs 1 loss; approximately 2.12 Mb total CNV span). On the other hand, South American (SA) isolates showed only deletions with respect to the reference (0 gains vs 57 losses; approximately 0.87 Mb) (**Figure 3B**). No gene displayed opposing CNV states (gain vs loss) between the two lineages, indicating non-overlapping targets of structural variation.

When comparing the functional repertoires of *Sporothrix brasiliensis* and *S. schenckii*, the two species showed dissimilar trends in gene gain and loss driven by CNV events. In *S. brasiliensis*, CNV-associated gains are mainly associated with molecular functions related to protein kinase activity, phosphotransferase activity, ATP binding, and proteins associated with microtubule trafficking (**Figure S4, Table S3**). Conversely, the losses in *S. brasiliensis* primarily involve genes encoding ribosomal proteins, structural constituents of translation, oxidoreductases, and enzymes involved in amino acid and lipid metabolism. (**Figure S4, Table S3**). For *S. schenckii,* the gains were focused on transporters, ion-binding proteins, and oxidoreductase enzymes. The losses included kinases and regulatory proteins, (**Figure S5, Table S4**). These findings indicate that the genes and gene categories affected by CNVs differ between both *Sporothrix* species.

### Genome-wide associations reveal multifactorial basis of itraconazole resistance in Sporothrix brasiliensis

Finally, a genome-wide association study (GWAS) was conducted to identify variants associated with resistance to itraconazole. A total of 81 SNPs that were identified genome-wide as significantly associated with the resistance trait (p < 1* 10^-5^); (**Figure 4; Table S5**). The analysis revealed single scattered SNPs rather than distinct genomic regions linked to the trait. Significant variants were distributed across multiple scaffolds. Among these, 29 SNPs mapped to coding regions, including 14 synonymous (SYN) and 15 nonsynonymous (NSY) substitutions, 50 to intergenic intervals (including upstream and downstream regulatory regions), and 2 in intronic region. The coding variants comprised of both nonsynonymous and synonymous substitutions (Table S5). Gene-level mapping highlighted several candidate genes, including SPBR_02168 (2 sites) and SPBR_00021, SPBR_00237, SPBR_00359, SPBR_00667, SPBR_02233, SPBR_02392, SPBR_02543, SPBR_03592, SPBR_03640, SPBR_04097, SPBR_05759, SPBR_05765, SPBR_06183, and SPBR_06651 (1 site each). All loci were subsequently annotated with the closest orthologs (Table S6). None of the alleles have been previously implicated in azole resistance. Gene Ontology (GO) analyses indicated that this dataset is enriched for ABC transporters (Genes=16, Fold enrichment=6.9; Enrichment FDR= 6.4 × 10^-11^), mismatch repair (Genes=9, Fold enrichment=3.5; Enrichment FDR= 5.6 × 10^-3^), and DNA repair (Genes=12, Fold enrichment=3.0; Enrichment FDR= 4.0 × 10^-3^).

**Figure 4.**
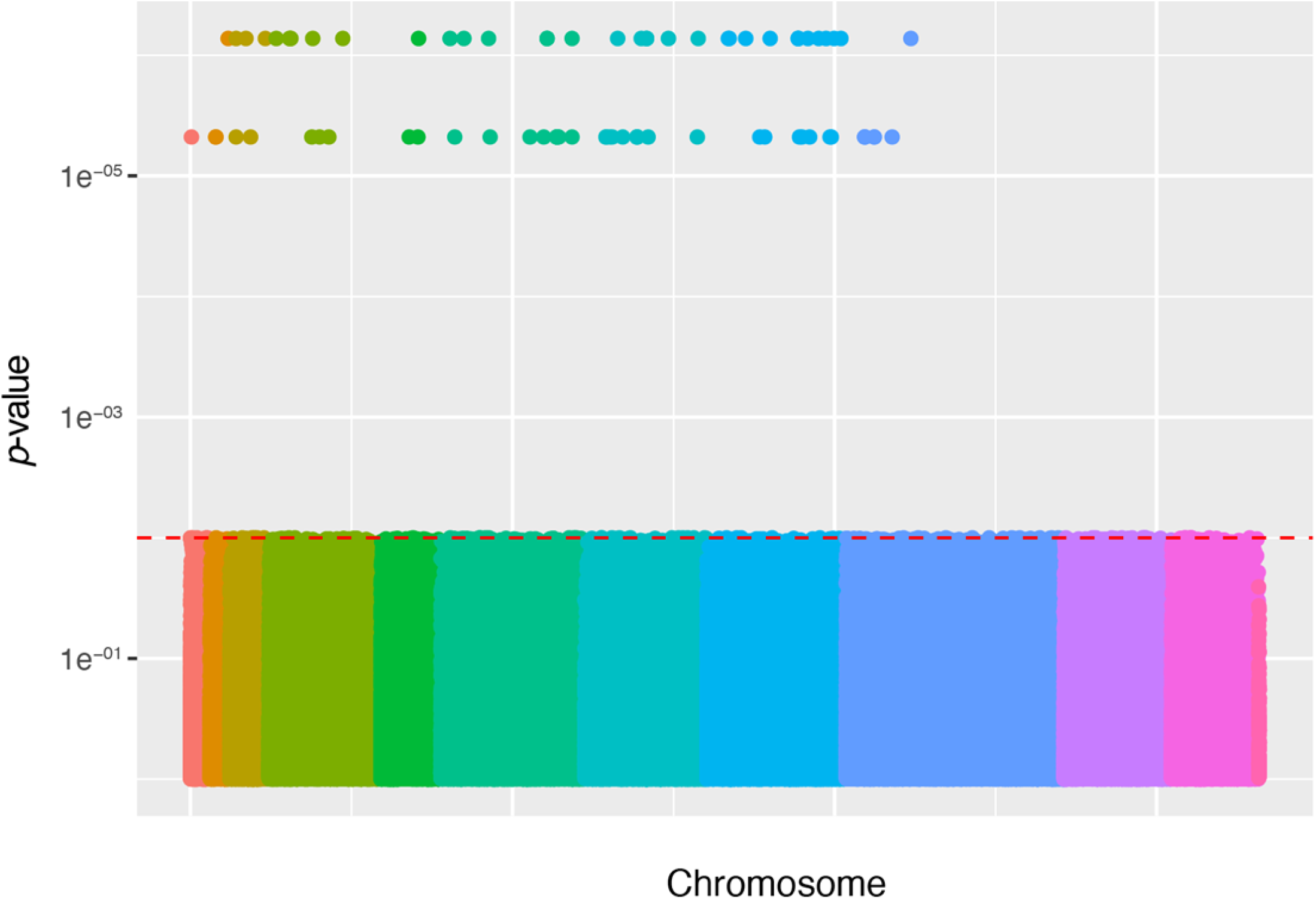
Genome-wide association analysis of itraconazole resistance in *Sporothrix brasiliensis*. Manhattan plot depicting the genome-wide distribution of SNPs associated with itraconazole resistance across all contigs. Each point represents an individual SNP, and colors alternate by chromosome for visual distinction. The y-axis shows the –log₁₀(*p*-value) of association under a binary resistance phenotype (MIC ≥ 8 µg/mL).

## Discussion

The magnitude and types of genetic variation in fungal pathogens represent an active area of research [44]. Deciphering this diversity is essential to identify the signatures of natural selection acting on pathogenic traits, including virulence, host adaptation, and antifungal tolerance. This question is particularly pressing for *Sporothrix*, a major One Health pathogen, where clonal expansion and geographic spread are reshaping disease burden but the evolutionary drivers of emergence are still unclear [2]. In this study, nucleotide and structural variation of *S. brasiliensis* and *S. schenckii* across the American continent were quantified. By combining SNP-based phylogenomics, CNV profiling, and GWAS, patterns of population structure, structural variation, and antifungal resistance in fungal emerging pathogens were uncovered. Our analyses indicate that *S. schenckii* exhibits substantially greater genomic diversity than *S. brasiliensis* [4, 23]. The deep phylogeographic split between the North and South American *S. schenckii* lineages, coupled with strong population structure suggests long-term geographic isolation (**Figure 1**). In contrast, *S. brasiliensis* clades are dominated by low diversity and uneven mating type distribution, which is indicative of rapid clonal expansion from a recent common ancestor [4, 10, 45]. The predominance of Clade Sb A across multiple and distant Brazilian states and hosts underscores the widespread dissemination of a successful epidemic genotype, while the existence of geographically restricted lineages (e.g., Clades C, D and E) reflects less sampled lineages that require further investigation [4, 10, 15, 46]. Across *S. brasiliensis* populations, mating-type structure was highly skewed, with most clades showing complete fixation for a single idiomorph **(Table I)**, consistent with predominantly clonal propagation. Notably, the Federal District (Clade D) was the only population in which both *MAT1-1* and *MAT1-2* were detected at equal frequency (1:1), indicating the coexistence of both mating types in this region [23]. The dominance of a single mating type across geographic clades supports multiple independent clonal emergences and dissemination events, consistent with founder effects and rapid regional spread driven largely by clonal transmission [4, 10, 15]

Evidence of admixed genotypes in certain *S. brasiliensis* isolates challenges the classical view of strict clonality and suggests the possibility of occasional genetic exchange [47, 48]. Although mosaic ancestry appears uncommon, its presence raises important questions about the degree of connectivity among *S. brasiliensis* lineages across their broad geographic distribution (**Figure 1**)[49]. Similar signals of recombination and intraspecific horizontal transfer have been reported in other fungal pathogens lacking a known sexual stage, indicating that genetic exchange can occur even when classical sexual reproduction has not been observed [50, 51]. Nonetheless, the underlying mechanism(s) generating these patterns in pathogenic *Sporothrix* (sexual, parasexual, or driven by other processes) remain unresolved despite decades of investigation [10, 52].

Our CNV analyses revealed different genomic patterns in the two *Sporothrix* species studied. *S. brasiliensis* showed a larger number of genes with evidence of CNVs (158 versus 88 in *S. schenckii*), with most events representing deletions rather than duplications. These CNVs were more prevalent in the subtelomeric regions **(Figure 2)**, which are recognized hotspots for adaptive evolution in pathogenic fungi [53], and preferentially affected genes involved in translation, metabolism, and cell wall biosynthesis. Conversely, CNV gains in *S. brasiliensis* were concentrated in genes related to kinase signaling, Golgi organization, and vesicle trafficking. *S. schenckii*, on the other hand, showed gains in transporters, oxidoreductases, and metabolic enzymes, while losses were primarily in regulatory or signaling elements. The underlying reasons of these differences might be ecological, demographic, or related to variations in sample size. A more systematic sampling approach, coupled with highly contiguous assemblies of both species, will be necessary to investigate these potential causes further.

The results presented here provide an initial approximation for identifying the genetic basis of azole resistance in *S. brasiliensis*. We identified 81 sites associated with high resistance to fluconazole scattered throughout the genome **(Figure 4)**. Notably, none of the alleles significantly associated with this trait have been previously implicated with azole resistance in any fungi. While it is possible that the genetic basis of azole resistance in *Sporothrix* may lead to the identification of novel molecular mechanisms, it is also possible that our analyses lack sufficient power to disentangle the genetic basis of this trait. This limitation is particularly important given that the number of isolates with standardized MIC data remains limited and non-standardized. Such constraints, combined with the population structure within *S. brasiliensis* complicate the interpretation of genetic mapping.

Our study has other caveats that warrant attention. First, the geographic sampling of *Sporothrix* remains uneven, with many endemic regions across the geographic range of the genus either underrepresented or unsampled. This uneven sampling may bias population-level inferences and underestimating the level of genetic diversity [54]. A coordinated network of genomic epidemiology is needed to understand the emergence of these pathogens [2]. Second, our detection of CNVs is promising but requires confirmation through new assemblies using long-read sequencing technology (see [55]). Additionally, the functional consequences of the variants detected in this study, identified through population genetic approaches, await evaluation within a functional genetics framework to establish causal links between candidate loci and resistance phenotypes.

The integration of comparative genomics, population structure, and association mapping establishes a foundational framework for dissecting the molecular basis of adaptation and drug resistance in *Sporothrix* [12]. Future work that expands the sample size, standardizes the MIC testing across reference centers and incorporating transcriptomic and metabolomic data will be crucial for translating these genomic insights into mechanistic understanding and clinical application [2, 56]. Ultimately, this study positions *S. brasiliensis* as a model for understanding recently emergent zoonotic pathogens. By unraveling the interplay between structural variation, selection, and drug response, we gain not only a clearer view of *Sporothrix* evolution but also a framework for monitoring antifungal resistance in other emerging fungal pathogens.

## ACKNOWLEDGMENTS

M.S.S.F., L.G.B.A., L.R.C., and M.M.T are supported by the Conselho Nacional de Desenvolvimento Científico e Tecnológico (CNPq) under grant numbers 409078/2022-0 and 445376/2023-6.

---

**Figure S1. Scaffold-based phylogenetic analyses reveal mosaic topologies within *Sporothrix brasiliensis*.** Maximum-likelihood (ML) phylogenies reconstructed from individual scaffolds of *S. brasiliensis* and *S. schenckii* show overall concordance with the whole-genome SNP tree but highlight local topological shifts among several *S. brasiliensis* isolates. Isolates CBS130107, CBS133004, CBS132985, and CBS132988 exhibit variable placement across scaffolds. Branch support values are derived from ultrafast bootstrap (UFBoot) and aBayes metrics.

**Figure S2. Chromosome-level distribution of copy number variants (CNVs) in *Sporothrix brasiliensis*.** Bar plots showing CNV counts for each chromosome across individual isolates of *S. brasiliensis*. Red bars represent copy number gains, and blue bars represent losses.

**Figure S3. Chromosome-level distribution of copy number variants (CNVs) in *Sporothrix schenckii*.** Bar plots showing CNV counts for each chromosome across individual isolates of *S. brasiliensis*. Red bars represent copy number gains, and blue bars represent losses.

**Figure S4.** Cytoscape bubble charts summarizing GO enrichment for CNV-associated genes in *Sporothrix brasiliensis*. Panels: (A) Biological Process—gains; (B) Molecular Function—gains; (C) Biological Process—losses; (D) Molecular Function—losses. Each bubble represents a GO term; bubble color encodes the Value provided with each GO term in the dataset, and bubble size reflects the term’s LogSize. Terms were summarized and visualized in Cytoscape to highlight functional themes among CNV gains versus losses.

**Figure S5.** Cytoscape bubble charts summarizing GO enrichment for CNV-associated genes in *Sporothrix schenkii*. Panels: (A) Biological Process—gains; (B) Molecular Function—gains; (C) Biological Process—losses; (D) Molecular Function—losses. Each bubble represents a GO term; bubble color encodes the Value provided with each GO term in the dataset, and bubble size reflects the term’s LogSize. Terms were summarized and visualized in Cytoscape to highlight functional themes among CNV gains versus losses.

## References

1. Denning DW. Global incidence and mortality of severe fungal disease. Lancet Infect Dis. 2024. Epub 2024/01/16. doi: 10.1016/S1473-3099(23)00692-8. PubMed PMID: 38224705.

2. Matute DR, Teixeira MM. Sporothrix is neglected among the neglected. PLoS Pathog. 2025;21(3):e1012898. Epub 20250306. doi: 10.1371/journal.ppat.1012898. PubMed PMID: 40048686; PubMed Central PMCID: PMCPMC11884826.

3. de Beer ZW, Duong TA, Wingfield MJ. The divorce of Sporothrix and Ophiostoma: solution to a problematic relationship. Studies in Mycology. 2016;83(1):165–91. doi: 10.1016/j.simyco.2016.07.001.

4. Zhang Y, Hagen F, Stielow B, Rodrigues AM, Samerpitak K, Zhou X, et al. Phylogeography and evolutionary patterns in Sporothrix spanning more than 14 000 human and animal case reports. Persoonia. 2015;35:1–20. doi: 10.3767/003158515X687416. PubMed PMID: 26823625; PubMed Central PMCID: PMCPMC4713101.

5. Santos MT, Nascimento LFJ, Barbosa AAT, Martins MP, Tunon GIL, Santos POM, et al. The rising incidence of feline and cat-transmitted sporotrichosis in Latin America. Zoonoses Public Health. 2024;71(6):609–19. Epub 20240723. doi: 10.1111/zph.13169. PubMed PMID: 39044549.

6. Hennessee I, Barber E, Petro E, Lindemann S, Buss B, Santos A, et al. Sporotrichosis Cluster in Domestic Cats and Veterinary Technician, Kansas, USA, 2022. Emerg Infect Dis. 2024;30(5):1053–5. doi: 10.3201/eid3005.231563. PubMed PMID: 38666748; PubMed Central PMCID: PMCPMC11060436.

7. Siew HH. The Current Status of Feline Sporotrichosis in Malaysia. Med Mycol J. 2017;58(3):E107–E13. doi: 10.3314/mmj.17.014. PubMed PMID: 28855477.

8. Orofino-Costa R, Freitas DFS, Bernardes-Engemann AR, Rodrigues AM, Talhari C, Ferraz CE, et al. Human sporotrichosis: recommendations from the Brazilian Society of Dermatology for the clinical, diagnostic and therapeutic management. An Bras Dermatol. 2022;97(6):757–77. Epub 20220922. doi: 10.1016/j.abd.2022.07.001. PubMed PMID: 36155712; PubMed Central PMCID: PMCPMC9582924.

9. Sanchotene KO, Madrid IM, Klafke GB, Bergamashi M, Della Terra PP, Rodrigues AM, et al. Sporothrix brasiliensis outbreaks and the rapid emergence of feline sporotrichosis. Mycoses. 2015;58(11):652–8. Epub 2015/09/26. doi: 10.1111/myc.12414. PubMed PMID: 26404561.

10. Teixeira Mde M, Rodrigues AM, Tsui CK, de Almeida LG, Van Diepeningen AD, van den Ende BG, et al. Asexual propagation of a virulent clone complex in a human and feline outbreak of sporotrichosis. Eukaryot Cell. 2015;14(2):158–69. doi: 10.1128/EC.00153-14. PubMed PMID: 25480940; PubMed Central PMCID: PMCPMC4311920.

11. Teixeira MM, de Almeida LG, Kubitschek-Barreira P, Alves FL, Kioshima ES, Abadio AK, et al. Comparative genomics of the major fungal agents of human and animal Sporotrichosis: Sporothrix schenckii and Sporothrix brasiliensis. BMC Genomics. 2014;15:943. doi: 10.1186/1471-2164-15-943. PubMed PMID: 25351875; PubMed Central PMCID: PMCPMC4226871.

12. Teixeira MM, Almeida-Paes R, Bernardes-Engemann AR, Nicola AM, de Macedo PM, Valle ACF, et al. Single nucleotide polymorphisms and chromosomal copy number variation may impact the Sporothrix brasiliensis antifungal susceptibility and sporotrichosis clinical outcomes. Fungal Genet Biol. 2022;163:103743. Epub 2022/09/25. doi: 10.1016/j.fgb.2022.103743. PubMed PMID: 36152775.

13. Ribeiro dos Santos A, Misas E, Min B, Le N, Bagal UR, Parnell LA, et al. Emergence of zoonotic sporotrichosis in Brazil: a genomic epidemiology study. The Lancet Microbe. 2024;5(3):e282–e90. doi: 10.1016/S2666-5247(23)00364-6.

14. de Souza Rabello VB, de Melo Teixeira M, Meyer W, Irinyi L, Xavier MO, Poester VR, et al. Multi-locus sequencing typing reveals geographically related intraspecies variability of Sporothrix brasiliensis. Fungal Genet Biol. 2024;170:103845. Epub 20231129. doi: 10.1016/j.fgb.2023.103845. PubMed PMID: 38040325.

15. de Carvalho JA, Hagen F, Fisher MC, de Camargo ZP, Rodrigues AM. Genome-wide mapping using new AFLP markers to explore intraspecific variation among pathogenic Sporothrix species. PLoS Negl Trop Dis. 2020;14(7):e0008330. Epub 2020/07/02. doi: 10.1371/journal.pntd.0008330. PubMed PMID: 32609739; PubMed Central PMCID: PMCPMC7329091.

16. Rodrigues AM, de Hoog G, Zhang Y, de Camargo ZP. Emerging sporotrichosis is driven by clonal and recombinant Sporothrix species. Emerg Microbes Infect. 2014;3(5):e32. Epub 2015/06/04. doi: 10.1038/emi.2014.33. PubMed PMID: 26038739; PubMed Central PMCID: PMCPMC4051365.

17. Almeida-Silva F, Coelho RA, Bernardes-Engemann AR, Fichman V, Freitas DF, Galhardo MC, et al. In vitro isavuconazole activity against Sporothrix brasiliensis suggests its efficacy in some severe sporotrichosis cases. Future Microbiol. 2023;18:1041–8. Epub 20230918. doi: 10.2217/fmb-2023-0094. PubMed PMID: 37721514.

18. Almeida-Paes R, Oliveira MME, Freitas DFS, Valle A, Gutierrez-Galhardo MC, Zancope-Oliveira RM. Refractory sporotrichosis due to Sporothrix brasiliensis in humans appears to be unrelated to in vivo resistance. Med Mycol. 2017;55(5):507–17. Epub 2016/10/25. doi: 10.1093/mmy/myw103. PubMed PMID: 27771622.

19. Espinel-Ingroff A, Abreu DPB, Almeida-Paes R, Brilhante RSN, Chakrabarti A, Chowdhary A, et al. Multicenter, International Study of MIC/MEC Distributions for Definition of Epidemiological Cutoff Values for Sporothrix Species Identified by Molecular Methods. Antimicrob Agents Chemother. 2017;61(10). Epub 2017/07/26. doi: 10.1128/AAC.01057-17. PubMed PMID: 28739796; PubMed Central PMCID: PMCPMC5610517.

20. Almeida-Paes R, Figueiredo-Carvalho MH, Brito-Santos F, Almeida-Silva F, Oliveira MM, Zancope-Oliveira RM. Melanins Protect Sporothrix brasiliensis and Sporothrix schenckii from the Antifungal Effects of Terbinafine. PLoS One. 2016;11(3):e0152796. doi: 10.1371/journal.pone.0152796. PubMed PMID: 27031728; PubMed Central PMCID: PMCPMC4816517.

21. Ribeiro Dos Santos A, Gade L, Misas E, Litvintseva AP, Nunnally NS, Parnell LA, et al. Bimodal distribution of azole susceptibility in Sporothrix brasiliensis isolates in Brazil. Antimicrob Agents Chemother. 2024;68(4):e0162023. Epub 20240222. doi: 10.1128/aac.01620-23. PubMed PMID: 38385701; PubMed Central PMCID: PMCPMC10989022.

22. Ferrer C, Colom F, Frases S, Mulet E, Abad JL, Alio JL. Detection and identification of fungal pathogens by PCR and by ITS2 and 5.8S ribosomal DNA typing in ocular infections. J Clin Microbiol. 2001;39(8):2873–9. doi: 10.1128/JCM.39.8.2873-2879.2001. PubMed PMID: 11474006; PubMed Central PMCID: PMCPMC88253.

23. Eudes Filho J, Santos IBD, Reis CMS, Patane JSL, Paredes V, Bernardes J, et al. A novel Sporothrix brasiliensis genomic variant in Midwestern Brazil: evidence for an older and wider sporotrichosis epidemic. Emerg Microbes Infect. 2020;9(1):2515–25. Epub 2020/11/07. doi: 10.1080/22221751.2020.1847001. PubMed PMID: 33155518; PubMed Central PMCID: PMCPMC7717857.

24. Sahl JW, Lemmer D, Travis J, Schupp JM, Gillece JD, Aziz M, et al. NASP: an accurate, rapid method for the identification of SNPs in WGS datasets that supports flexible input and output formats. Microb Genom. 2016;2(8):e000074. Epub 20160825. doi: 10.1099/mgen.0.000074. PubMed PMID: 28348869; PubMed Central PMCID: PMCPMC5320593.

25. Bolger AM, Lohse M, Usadel B. Trimmomatic: a flexible trimmer for Illumina sequence data. Bioinformatics. 2014;30(15):2114–20. doi: 10.1093/bioinformatics/btu170. PubMed PMID: 24695404; PubMed Central PMCID: PMCPMC4103590.

26. Danecek P, Bonfield JK, Liddle J, Marshall J, Ohan V, Pollard MO, et al. Twelve years of SAMtools and BCFtools. Gigascience. 2021;10(2). Epub 2021/02/17. doi: 10.1093/gigascience/giab008. PubMed PMID: 33590861; PubMed Central PMCID: PMCPMC7931819.

27. Kurtz S, Phillippy A, Delcher AL, Smoot M, Shumway M, Antonescu C, Salzberg SL. Versatile and open software for comparing large genomes. Genome Biol. 2004;5(2):R12. Epub 2004/02/05. doi: 10.1186/gb-2004-5-2-r12. PubMed PMID: 14759262; PubMed Central PMCID: PMCPMC395750.

28. Delcher AL, Phillippy A, Carlton J, Salzberg SL. Fast algorithms for large-scale genome alignment and comparison. Nucleic Acids Res. 2002;30(11):2478–83. Epub 2002/05/30. doi: 10.1093/nar/30.11.2478. PubMed PMID: 12034836; PubMed Central PMCID: PMCPMC117189.

29. Li H, Durbin R. Fast and accurate short read alignment with Burrows-Wheeler transform. Bioinformatics. 2009;25(14):1754–60. doi: 10.1093/bioinformatics/btp324. PubMed PMID: 19451168; PubMed Central PMCID: PMCPMC2705234.

30. McKenna A, Hanna M, Banks E, Sivachenko A, Cibulskis K, Kernytsky A, et al. The Genome Analysis Toolkit: a MapReduce framework for analyzing next-generation DNA sequencing data. Genome Res. 2010;20(9):1297–303. Epub 20100719. doi: 10.1101/gr.107524.110. PubMed PMID: 20644199; PubMed Central PMCID: PMCPMC2928508.

31. Li H. Minimap2: pairwise alignment for nucleotide sequences. Bioinformatics. 2018;34(18):3094–100. Epub 2018/05/12. doi: 10.1093/bioinformatics/bty191. PubMed PMID: 29750242; PubMed Central PMCID: PMCPMC6137996.

32. Minh BQ, Schmidt HA, Chernomor O, Schrempf D, Woodhams MD, von Haeseler A, Lanfear R. IQ-TREE 2: New Models and Efficient Methods for Phylogenetic Inference in the Genomic Era. Mol Biol Evol. 2020;37(5):1530–4. Epub 2020/02/06. doi: 10.1093/molbev/msaa015. PubMed PMID: 32011700; PubMed Central PMCID: PMCPMC7182206.

33. Kalyaanamoorthy S, Minh BQ, Wong TKF, von Haeseler A, Jermiin LS. ModelFinder: fast model selection for accurate phylogenetic estimates. Nat Methods. 2017;14(6):587–9. doi: 10.1038/nmeth.4285. PubMed PMID: 28481363; PubMed Central PMCID: PMCPMC5453245.

34. Minh BQ, Nguyen MA, von Haeseler A. Ultrafast approximation for phylogenetic bootstrap. Mol Biol Evol. 2013;30(5):1188–95. doi: 10.1093/molbev/mst024. PubMed PMID: 23418397; PubMed Central PMCID: PMCPMC3670741.

35. Anisimova M, Gascuel O. Approximate likelihood-ratio test for branches: A fast, accurate, and powerful alternative. Syst Biol. 2006;55(4):539–52. Epub 2006/06/21. doi: 10.1080/10635150600755453. PubMed PMID: 16785212.

36. Anisimova M, Gil M, Dufayard JF, Dessimoz C, Gascuel O. Survey of branch support methods demonstrates accuracy, power, and robustness of fast likelihood-based approximation schemes. Syst Biol. 2011;60(5):685–99. Epub 20110503. doi: 10.1093/sysbio/syr041. PubMed PMID: 21540409; PubMed Central PMCID: PMCPMC3158332.

37. Raj A, Stephens M, Pritchard JK. fastSTRUCTURE: variational inference of population structure in large SNP data sets. Genetics. 2014;197(2):573–89. doi: 10.1534/genetics.114.164350. PubMed PMID: 24700103; PubMed Central PMCID: PMCPMC4063916.

38. Alexander DH, Novembre J, Lange K. Fast model-based estimation of ancestry in unrelated individuals. Genome Res. 2009;19(9):1655–64. Epub 2009/08/04. doi: 10.1101/gr.094052.109. PubMed PMID: 19648217; PubMed Central PMCID: PMCPMC2752134.

39. Purcell S, Neale B, Todd-Brown K, Thomas L, Ferreira MA, Bender D, et al. PLINK: a tool set for whole-genome association and population-based linkage analyses. Am J Hum Genet. 2007;81(3):559–75. doi: 10.1086/519795. PubMed PMID: 17701901; PubMed Central PMCID: PMCPMC1950838.

40. Brown MR, Manuel Gonzalez de La, Rosa P, Blaxter M. tidk: a toolkit to rapidly identify telomeric repeats from genomic datasets. Bioinformatics. 2025;41(2):btaf049. doi: 10.1093/bioinformatics/btaf049.

41. Su W, Ou S, Hufford MB, Peterson T. A Tutorial of EDTA: Extensive De Novo TE Annotator. Methods Mol Biol. 2021;2250:55–67. doi: 10.1007/978-1-0716-1134-0_4. PubMed PMID: 33900591.

42. Boeva V, Popova T, Bleakley K, Chiche P, Cappo J, Schleiermacher G, et al. Control-FREEC: a tool for assessing copy number and allelic content using next-generation sequencing data. Bioinformatics. 2012;28(3):423–5. Epub 20111206. doi: 10.1093/bioinformatics/btr670. PubMed PMID: 22155870; PubMed Central PMCID: PMCPMC3268243.

43. Foundation PS. Python. 3.9.13 ed2024.

44. Fisher MC, Gurr SJ, Cuomo CA, Blehert DS, Jin H, Stukenbrock EH, et al. Threats Posed by the Fungal Kingdom to Humans, Wildlife, and Agriculture. mBio. 2020;11(3). Epub 2020/05/07. doi: 10.1128/mBio.00449-20. PubMed PMID: 32371596; PubMed Central PMCID: PMCPMC7403777.

45. Rodrigues AM, de Melo Teixeira M, de Hoog GS, Schubach TM, Pereira SA, Fernandes GF, et al. Phylogenetic analysis reveals a high prevalence of Sporothrix brasiliensis in feline sporotrichosis outbreaks. PLoS Negl Trop Dis. 2013;7(6):e2281. doi: 10.1371/journal.pntd.0002281. PubMed PMID: 23818999; PubMed Central PMCID: PMCPMC3688539.

46. Rodrigues AM, de Hoog GS, de Camargo ZP. Sporothrix Species Causing Outbreaks in Animals and Humans Driven by Animal-Animal Transmission. PLoS Pathog. 2016;12(7):e1005638. Epub 2016/07/16. doi: 10.1371/journal.ppat.1005638. PubMed PMID: 27415796; PubMed Central PMCID: PMCPMC4945023.

47. Matute DR, Sepulveda VE. Fungal species boundaries in the genomics era. Fungal Genet Biol. 2019;131:103249. Epub 2019/07/08. doi: 10.1016/j.fgb.2019.103249. PubMed PMID: 31279976.

48. Sasaki AA, Fernandes GF, Rodrigues AM, Lima FM, Marini MM, Dos SFL, et al. Chromosomal polymorphism in the Sporothrix schenckii complex. PLoS One. 2014;9(1):e86819. doi: 10.1371/journal.pone.0086819. PubMed PMID: 24466257; PubMed Central PMCID: PMCPMC3900657.

49. Meier CS, Pagni M, Richard S, Mühlethaler K, Almeida JMGCF, Nevez G, et al. Fungal antigenic variation using mosaicism and reassortment of subtelomeric genes’ repertoires. Nature Communications. 2023;14(1):7026. doi: 10.1038/s41467-023-42685-6.

50. Habig M, Grasse AV, Müller J, Stukenbrock EH, Leitner H, Cremer S. Frequent horizontal chromosome transfer between asexual fungal insect pathogens. Proceedings of the National Academy of Sciences. 2024;121(11):e2316284121. doi: 10.1073/pnas.2316284121.

51. Taylor J, Jacobson D, Fisher M. THE EVOLUTION OF ASEXUAL FUNGI: Reproduction, Speciation and Classification. Annu Rev Phytopathol. 1999;37:197–246. doi: 10.1146/annurev.phyto.37.1.197. PubMed PMID: 11701822.

52. Fuchs T, Visagie Cobus M, Wingfield Brenda D, Wingfield Michael J. Sporothrix and Sporotrichosis: A South African Perspective on a Growing Global Health Threat. Mycoses. 2024;67(10):e13806. doi: 10.1111/myc.13806.

53. Vande Zande P, Zhou X, Selmecki A. The Dynamic Fungal Genome: Polyploidy, Aneuploidy and Copy Number Variation in Response to Stress. Annu Rev Microbiol. 2023;77:341–61. Epub 20230612. doi: 10.1146/annurev-micro-041320-112443. PubMed PMID: 37307856; PubMed Central PMCID: PMCPMC10599402.

54. Dewar AE, Belcher LJ, West SA. A phylogenetic approach to comparative genomics. Nature Reviews Genetics. 2025;26(6):395–405. doi: 10.1038/s41576-024-00803-0.

55. Spealman P, De T, Chuong JN, Gresham D. Best Practices in Microbial Experimental Evolution: Using Reporters and Long-Read Sequencing to Identify Copy Number Variation in Experimental Evolution. J Mol Evol. 2023;91(3):356–68. Epub 20230403. doi: 10.1007/s00239-023-10102-7. PubMed PMID: 37012421; PubMed Central PMCID: PMCPMC10275804.

56. Rossow JA, Queiroz-Telles F, Caceres DH, Beer KD, Jackson BR, Pereira JG, et al. A One Health Approach to Combatting Sporothrix brasiliensis: Narrative Review of an Emerging Zoonotic Fungal Pathogen in South America. J Fungi (Basel). 2020;6(4). Epub 2020/10/30. doi: 10.3390/jof6040247. PubMed PMID: 33114609; PubMed Central PMCID: PMCPMC7712324.

